# Ecological drivers of social complexity: the role of predation risk and nesting resource in group-living cichlid

**DOI:** 10.64898/2026.06.26.734653

**Authors:** Yuki Yoshio, Yoshimitsu Takada, Ryo Hidaka, Ryoichi Inoue, Kota Kambe, Shun Satoh

## Abstract

Understanding how social complexity responds to environmental variation remains a longstanding challenge in evolutionary biology. Here, we investigated the drivers of social complexity using intraspecific social variation across seven locations of the obligatory shell-brooding cichlid *Neolamprologus meeli* in Lake Tanganyika. We quantified the number of subordinate individuals per female territory and examined the effects of predation risk, shell availability, and their interaction. Social complexity increased with shell availability under high predation risk but showed little association under low predation risk. A field manipulative-experiment further demonstrated that increasing shell availability led to higher juvenile retention, indicating a causal effect of territory quality. In addition, removal of subordinates reduced shell availability, suggesting the feedback between group size and territory maintenance. We also assessed genetic population structure based on nuclear SNPs obtained by MIG-seq and found only weak genetic differentiation among localities, suggesting that the observed social variation is unlikely to simply reflect strong genetic subdivision. Together, these results show that predation risk promotes group living, whereas nesting resource availability constrains its extent. Our study highlights that social complexity emerges from the interaction between macro- and micro-ecological factors, providing a mechanistic understanding of the evolution of social complexity and philopatry.

## Introduction

Social complexity in organisms is highly diverse and often converges on similar forms across taxonomic groups. As a striking example of such complexity, cooperative breeding has long attracted the attention of evolutionary biologists. Cooperative breeding is generally defined as a social system in which non-breeding subordinate helpers delay dispersal, forego their own reproduction, and assist in caring for offspring that are not necessarily their own[1] . Such societies have been reported from various vertebrate taxa, such as mammals [2], birds [3,4], and fishes [5–7]. Traditionally, indirect fitness benefits arising through kin selection, which is meditated by genetic relatedness, have been considered as major evolutionary drivers of cooperative breeding system [2,3,8,9]. Although recent theoretical work suggests that high relatedness among group members can indeed promote the evolution of cooperative breeding [10], relatedness alone cannot fully account for why some individuals engage in cooperative breeding whereas conspecifics do not [11–13]. Understanding variation in cooperative breeding therefore requires moving beyond kin selection to consider direct fitness benefits, and to examine the factors that generate these benefits at both macro-ecological factors such as climatic conditions, and micro-ecological factors such as group composition and interactions among group members.

In birds, climatic conditions are widely recognized as one of the major macro-ecological factors promoting cooperative breeding. For example, in sociable weavers [14] and white-fronted bee-eaters [15], the number of helpers per nest is significantly higher in dry years than in wet years [14–16], and larger groups exhibit higher fledging success, with this effect being most pronounced under the harshest conditions [14,15,17]. As another example of macro-ecological factors, in the pied kingfisher, subordinate individuals are more likely to delay dispersal and form larger groups in populations where food resources are nutritionally poor and difficult to obtain [11,18].

Predation risk has been suggested to play a key role in facilitating delayed dispersal and social complexity among cichlid fish [19–21]. For group-living species in particular, predation profoundly influences social evolution: living in groups reduces predation risk through dilution [22,23], confusion, and collective anti-predator behaviors such as vigilance and mobbing [24]. Consequently, predation pressure is closely associated with group size, social behavior, and the evolution of cooperative systems [8,25]. Evidence from cooperatively breeding cichlid fishes supports this idea. In the Lake Tanganyika cichlids, *Neolamprologus pulcher* and *N. obscurus*, group composition are associated with the abundance of larger fish predators that particularly threaten smaller individuals [19]. In populations exposing higher predation pressure, subordinate individuals are more likely to delay dispersal and remain in their natal territories [19,20]. By remaining in their natal territory, subordinates can access not only to shelters [20] but also shared territory maintenance and group defense for predators [26,27]. This reciprocal exchange of benefits of breeders and subordinates is thought to facilitate the evolution of complex social systems in fish [21,28–31]. Consistent with this idea, recent phylogenetic analyses of Lake Tanganyika cichlids suggest that linages with smaller body size, which are exposed to greater predation risk, have repeatedly evolved cooperative breeding [7]. Therefore, predation risk may act as a macro-ecological factor that shapes dispersal decisions through interactions with micro-ecological factors, thereby promoting social complexity.

In shell brooding cichlids, suitable empty snail shells are essential for breeding and shelter resources. Because subordinate individuals depend on access to shells for survival and future reproduction, variation in shell availability is expected to influence the costs and benefits of remaining within social groups. Therefore, the obligated shell-dwelling cichlid, *N. meeli,* endemic to Lake Tanganyika, provides an ideal model system for testing how macro-and micro-ecological factors interact to shape social complexity. *Neolamprologus meeli* was described by Saeki et al. (2022) as a cooperatively breeding cichlid in which delayed-dispersing related subordinates engage in defense against intruders and nest maintenance [6]. Moreover, *N. meeli* belongs to a phylogenetically distinct clade from the other cooperatively breeding Lake Tanganyika cichlids that have been the main focus of previous studies [7]. This further highlights the value of this species as an independent model system for understanding the evolution of social complexity in Lake Tanganyika cichlids. Previous work has shown that delayed dispersal varies among breeding groups in *N. meeli,* with some nests containing subordinate individuals and others lacking them even within the same localities [6]. In the present study, we further show that social complexity, measured as the number of subordinates per nest, also varies markedly among localities (table 1).

**Table 1.** Variation in the social organization of *Neolamprologus meeli* among localities. The table presents the number of focal territories surveyed and the number of subordinates per territory at each locality. The number of subordinates is reported as the mean ± SE, median [interquartile range], and range. “Breeding females with subordinates” indicates the number and percentage of territories in which at least one subordinate was observed. BF, breeding female; sub., subordinate(s).

| locality | n territories | The number of sub. (mean±SE) | median [IQR] | Range | BF with sub.(%) |
| --- | --- | --- | --- | --- | --- |
| NK | 59 | 1.36 ± 0.154 | 1[0-2] | 0-4 | 41 (69.5%) |
| NKS | 38 | 0.421 ± 0.117 | 0[0-1] | 0-2 | 11 (28.9%) |
| CH | 27 | 0.667 ± 0.119 | 1[0-1] | 0-2 | 16 (59.3%) |
| MW | 21 | 0.000 | 0[0-0] | 0-0 | 0 (0.0%) |
| WZ | 58 | 0.655 ± 0.091 | 1[0-1] | 0-2 | 31 (53.4%) |
| KTS | 17 | 0.529 ± 0.194 | 0[0-1] | 0-2 | 6 (35.3%) |
| KTD | 9 | 0.000 | 0[0-0] | 0-0 | 0 (0.0%) |

Therefore, we investigated how ecological factors influence variation in social complexity (i.e. the number of subordinates per nest) across seven natural locations of *N.meeli* in Lake Tanganyika. Specifically, we examined the effect of predation risk, shell availability (as a measure of nesting resource abundance) and other ecological factors such as density of habitat competitors and food competitors. We also conducted behavioral observations in each locality to examine whether the major helping behaviors in *N. meeli*—defense against intruders and nest maintenance—were associated with social complexity, and whether the presence of subordinates influenced the behavior of breeding individuals. Moreover, to elucidate the relationship between social complexity and territory quality directly, we conducted experimental manipulations for the number of shells in breeding nests and subordinates. Because social variation among localities may also arise through genetic divergence, we additionally examined population genetic structure using nuclear SNPs data generated by MIG-seq.

## Materials and Methods

### Study species and fieldworks

*Neolamprologus meeli* is small substrate-brooding cichlid endemic to Lake Tanganyika. Although maximum body size in breeding males are 59.36 cm in standard length (SL) and breeding females are 52.86 cm SL [6], the body size are slightly differ among localities (see Results). Fieldwork was conducted at seven localities from the southern end of the lake between 2022 and 2025 using SCUBA diving. At Nkumbula Island, *N.meeli* occurs in both sandy-muddy bottoms (figure 1a) and shell beds (figure 1b); therefore, these were treated as separate localities (NK and NKS, respectively). At other localities (Wonzye; WZ, Chikonde; CH), individuals were found only on sandy-muddy substrates. In Mwina (MW), however, coarse gravel was spread on the bottom. In Katoto, sand was spread on the bottom and habitats were further divided into shallow (depth< 10 m; KTS) and deep (depth> 15 m; KTD) sites due to apparent difference in community composition.

**Figure 1.**
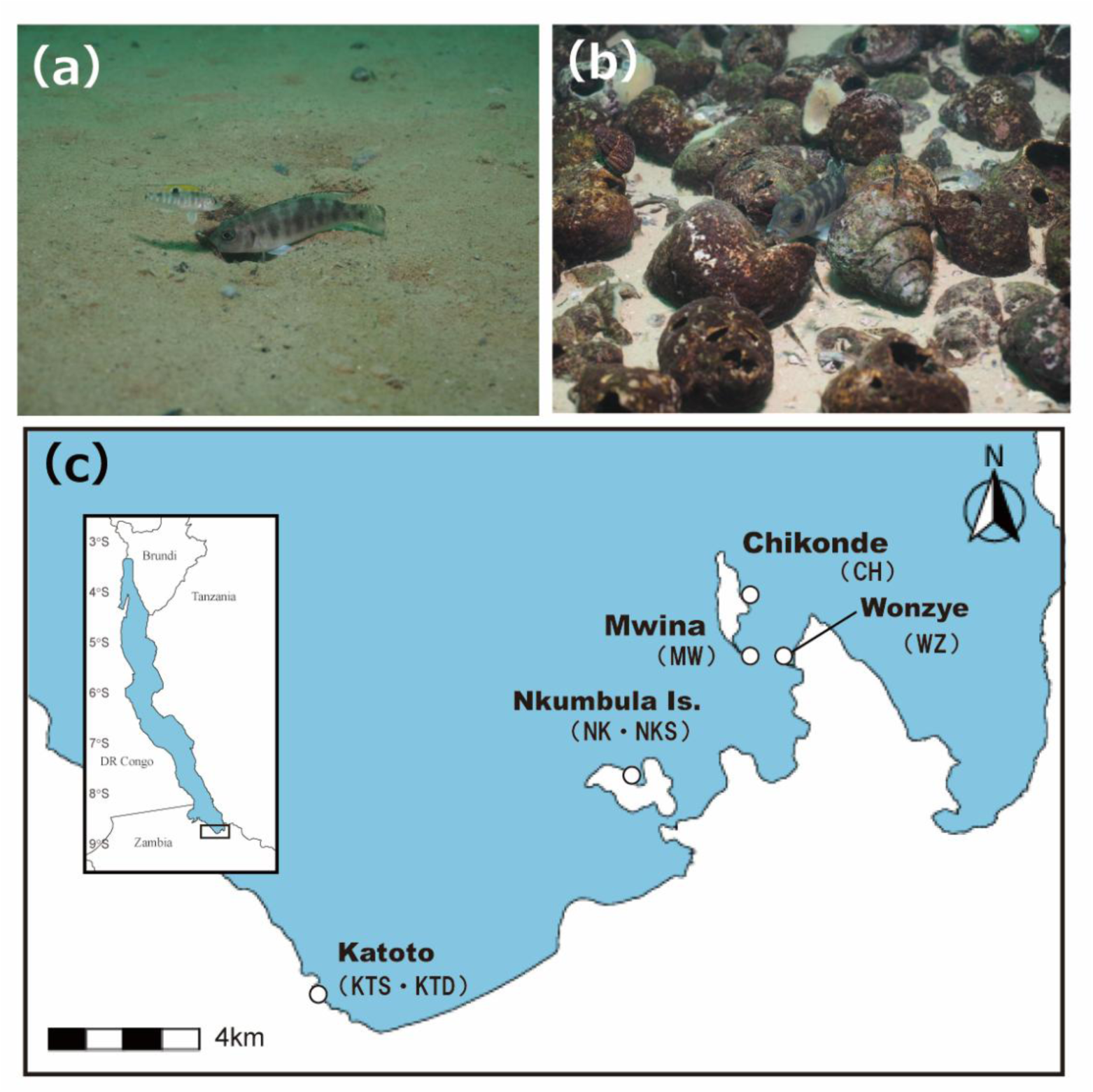
Study localities and breeding habitats of the shell-brooding cichlid *Neolamprologus meeli.* Representative breeding habitats on (a) a mixed sand–mud substrate and (b) a shell bed (photographs by S. S.). (c) Map of southern Lake Tanganyika showing the seven study localities.

### Definition of social status and composition

Individuals were classified into four social categories based on body size, behavior, and gonadal characteristics: breeding males, breeding females, subordinates, and juveniles. Breeding males were the largest individuals and exhibited wide-ranging movements across multiple territories. Breeding females were the second-largest individuals, occupied a single territory, and constructed crater-shaped nests. Subordinates were smaller individuals (>25.0 mm standard length, SL) that shared nests with breeding females and exhibited submissive behavior. These individuals also actively participated in helping activities, including territory defense and nest maintenance. Saeki et al. (2022), based on samples from Chikonde (CH; Mutondwe Island: 8°71.3′S, 31°12.3′W), defined subordinates as individuals larger than 30.0 mm SL [6]. However, because we observed locality-level variation in size at sexual maturity, we applied a lower threshold of 25.0 mm SL for classifying subordinates in this study and individuals smaller than 25.0 mm SL were classified as juveniles. The definitions of social status and the associated behavioral observations followed the methodology established in previous studies [6,32].

### Behavioral Observation and collecting data of nesting resource

We recorded behaviors of breeding females in each localities using underwater video cameras (GoPro Hero 11/12). Upon identifying a territory, we estimated the number of fish around it for about 5 min. Cameras positioned overhead, with 50×50 cm quadrat placed to substantially enclose for estimating SL and recording the number of available shells within the quadrat. Available shells are empty shells which aperture are not buried by sediment. Fifteen-minute video recordings were conducted once per nest, in the morning (9:00 - 11:00) or afternoon (13:00 - 15:00). For behavioral analysis, we recorded the frequency of aggressive behaviors against intruders (frontal display or overt aggression) and maintenance behaviors (digging and removing sand or removal of debris from the territory) as described in *N.meeli* [6]. All observations were restricted to individuals within a 50 × 50 cm quadrat centered on the nest.

### Sampling and measurements

We analyzed 319 individuals across all localities (table 2). Individuals were captured using gill nets and hand nets, transported to a field laboratory, and euthanized with an overdose of anesthetic (FA100, 10% eugenol solution). Gonadal inspection was used to confirm sex whenever possible, particularly for breeding females. Subordinates and juveniles were not sexed because their gonads were not detectable upon dissection. Specimens were fixed in 10% formalin and preserved in 70% ethanol. Standard length (SL) was measured to the nearest 0.01 mm. Fin clips were preserved in 100% ethanol. A total of 42 specimens from six area (NK:7, CH:8, MW:8, WZ:8, KTS:3, KTD:8) were used for genetic analysis. Because between NK and NKS are closely continued, we regard NK as same sites.

**Table 2.**
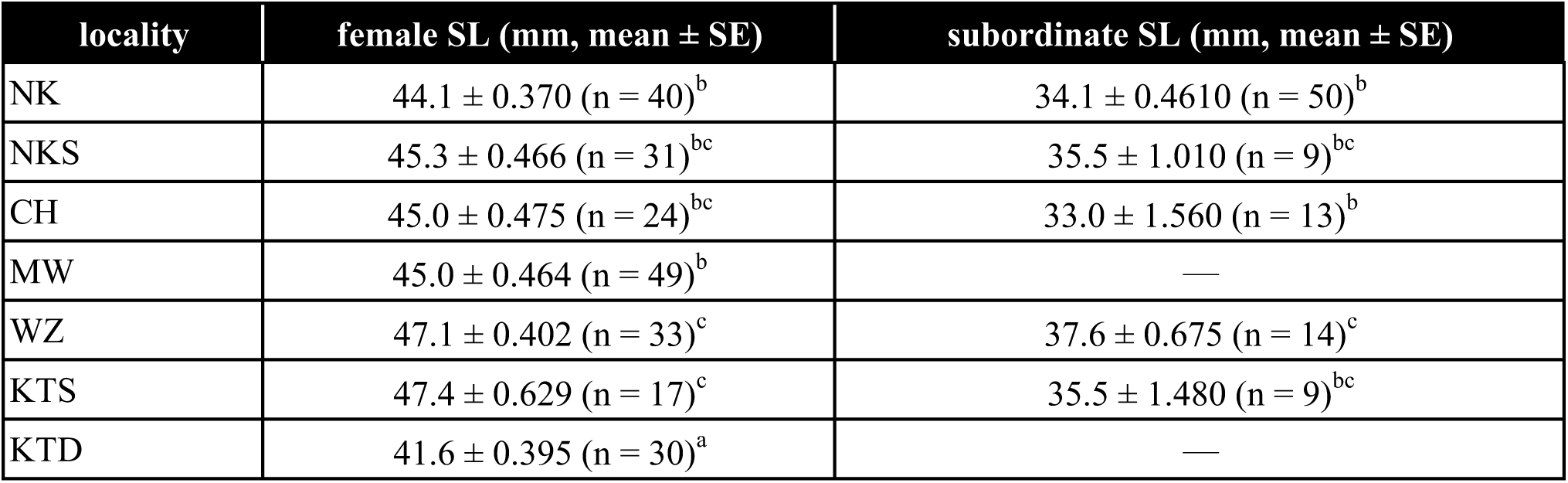
Differences among localities in the standard length (SL) of breeding females and subordinates. Values are presented as the mean ± SE. Different letters indicate significant differences among localities based on Tukey’s HSD test (p < 0.05).

### Habitat competitor and Food competitor

We estimated the density of habitat competitors in each locality by counting the number of habitat competitors that crossed 50 × 50 cm quadrats enclosing the crater-shaped nests o*f N. meeli* from videotaped data. We regarded habitat competitors as two shell-brooding cichlids, including *Telmatochromis temporalis* and *Lamprologus ocellatus*. Moreover, in only Mwina point, we regard *N. pulcher* as a habitat competitor because we observed *N. pulcher* also inhabits empty shells, similar to other shell-brooders. As same ways, we also counted the number of food competitors. It has been reported that *N. meeli* feeds invertebrate prey, such as arthropoda from substrate surface or into shell [6,33–35]. Therefore, food competitors were defined as cichlid fishes that feed on arthropods, including *Xenotilapia* spp., *Lamprologus callipterus*, *Gnathochromis lemairii*, and small piscivorous cichlids (< 8 cm SL), such as *Lepidiolamprologus elongatus*, *L. attenuatus*, *L. cunningtoni*, and *N. tetracanthus* [33,36,37]. We estimated the densities of habitat competitors and food competitors by randomly selecting a continuous 15-min segment from the video recording of each nest and counting the number of individuals that crossed the 50 × 50 cm quadrat during that period. These data were therefore obtained at the nest level. For locality-level analyses, we averaged the nest-level values within each population and used these means as locality-level estimates of habitat-competitor and food-competitor abundance.

### Estimation of the predation risk and scale-eating

From October to December 2025, we estimated predation risk in each locality by counting the number of piscivorous fish that attacked a decoy. Before each trial, we set up a decoy device underwater. One individual of *N.meeli* (mean±SE (33.8±1.91 cm) in standard length; *N. meeli*) was placed as a decoy in a transparent plastic bag (34 × 48 cm). A small amount of air was added so that the bag floated underwater. After the opening of the plastic bag was secured, the bag was fastened with a double clip. The length of the string was then adjusted so that the bag containing the decoy floated approximately 10–20 cm above the lake bottom. The decoy setup was enclosed within a 50 × 50 cm quadrat. We then placed a waterproof video camera (GoPro Hero 11/12) so that it recorded the decoy setup. The camera recorded for 1 h, and we immediately moved away from the setup after starting the recording to minimize any effect of diver presence on the fish community around the experimental area. We repeated these trials four to five times per locality on different days to capture temporal variation in fish activity. The fish used in the decoy device were replaced for each trial, and no fish were injured or died during the experiment. In total, we conducted these trials 31 times (NK:5, NKS:5, CH:4, MW:4, WZ:5, KTS:3, KTD:5). We counted the number of large piscivorous cichlids (>8 cm) that attacked the decoy, including *Lepidiolamprologus elongatus*, *L. attenuatus*, *L. cunningtoni*, *Neolamprologus tetracanthus*, and *Bathybates* spp. [39,47,48]. For each locality, count data were first averaged within each day and then averaged across all experimental days to obtain an index of predation risk.

In addition, *Perissodus microlepis* specializes in scale eating [49] and can damage shell-dwelling cichlids through scale-feeding attacks. Therefore, we also estimated the abundance or attack frequency of *P. microlepis* in each locality using the same method.

### Experimental manipulation for shell availability in breeding nest

To examine how territory quality influences juvenile dispersal under high predation risk, we experimentally manipulated the availability of nesting resources, namely empty snail shells, in a natural locality. The experiment was conducted at the Nkumbula sandy–muddy bottom site (NK), where predation risk is relatively high (figure 1b; table S1). We selected breeding nests containing juveniles (< 25 mm SL) and enclosed each nest within a 50 × 50 cm quadrat. In the shell-addition treatment, we added empty snail shells until the total number of usable shells within each quadrat reached 20, including shells originally present in the territory. In the control treatment, shells were also temporarily added and then removed, leaving shell availability unchanged from the original level. This procedure controlled potential disturbance effects associated with shell manipulation. Ten days after the manipulation, we counted the number of juveniles remaining in each nest. We used this measure to assess how increased shell availability affected changes in juvenile number within the natal territory, reflecting both juvenile dispersal and recruitment. The experiment included 11 control nests and 11 shell-addition nests.

### Removal experiment of subordinates and effects on shell availability

We conducted a subordinate-removal experiment to examine changes in shell availability following the loss of subordinates from territories exposed to high predation pressure. The experiment was conducted at the sandy–muddy bottom site at Nkumbula (NK) in December 2025. We haphazardly selected focal nests containing at least one subordinate and identified the center of each breeding nest through brief behavioral observations. We then centered a 50 × 50 cm quadrat on each nest and recorded the numbers of subordinates and available shells within the quadrat. All subordinates were subsequently removed, leaving only the breeding individuals. Because *N. meeli* inhabits sandy substrates, exposed shells may gradually become buried when nest maintenance declines. To quantify temporal changes in shell availability following subordinate removal, we counted the available shells within each quadrat immediately after removal and again four days later. We calculated the change in shell availability as the number recorded four days after removal minus the number recorded immediately after removal. This experiment allowed us to test whether shell availability declined during the four days following subordinate removal.

### Analysis of genetic data

To get nuclear SNP data, we used the multiplexed ISSR genotyping by sequencing method (MIG-seq[38]). Total genomic DNA was extracted from fin clips of 42 individuals from the Wizard Genomic DNA Purification Kit (Promega, Madison, WI, USA) (table S2). The MIG-seq library was prepared according to Onuki and Fuke (2020)[39]. The first round of PCR was conducted using eight ISSR primer sets with tail sequences (Onuki and Fuke 2022)[39] at an annealing temperature of 38 ℃. The Multiplex PCR Assay kit ver. 2 (Takara, Shiga) was used for the first PCR. The first PCR products were diluted and used in the second PCR. The second PCR was conducted using 24 forward and eight reverse primer pairs designed for the NovaSeq platform (Illumina[39]). Phusion High-Fidelity PCR Master Mix with HF Buffer (NEB, Ipswich, MA) was used for the second PCR, consisting of 20 cycles of 98 ℃ for 30s, 54℃ for 15s, and 68℃ for 30s. The products of the second PCR were mixed and purified usinga GeneRead Size Selection kit (Qiagen, Hilden). Fragment with size ranges of 300-800 bp were isolated and purified using SPRIselect (Beckman Coulter, Brea, CA). Fragment size and final concentration were confirmed using Agilent 2200 TapeStation (Beijing, China) and sequenced on Illumina NovaSeq X plus. The raw sequence data were deposited in the NCBI Sequence Read Archive (accession no. PRJNA1481448)

The raw data were demultiplexed using the process _shortbreads programs is Stacks v2.6 [40]. Quality control and trimming of the raw reads were performed using fastp v0.23.4 [41], during which primer sequences were trimmed, and low-quality bases (< Q30) as well as adapter sequences were removed. The trimmed reads were mapped to reference genome of *Neolamprologus multifasciatus* (accession no. ERS16590192) using bwa-mem2 v2.0pre2 [42] and the data files were converted to binary alignment map (BAM). To remove multi-hit reads, those containing XA:Z or SA:Z tags were excluded. The mapping rate was 57.38% on average (range: 47.85%-67.45%). The variant call was performed using the mpileup program in SAMtools v1.23.1 [43]. We filtered variant sites using the VCFtools v0.1.16 [44] with the following setting: --minQ 30, --remove-indels, --minDP 10, --min-meanDP 12, --maxDP 240, --max-alleles 2, --min-alleles 2, --mac 2 and --max-missing 0.9; the PLINK v1.90b7.7 [45]: -- indep-pairwise 50 10 0.25; and the populations program in Stacks: --write-single-snp, -R 0.7, --min-mac 2, --max-obs-het 0.75. After these procedures, 918 SNPs from 42 individuals were gotten. To estimate the population structure of *N. meeli*, we performed unsupervised clustering using the likelihood model-based program ADMIXTURE v1.3.0 [46] assuming the number of populations (K) from 1 to 7. These analyses were repeated 100 times for each K with different seed values, and the optimal K value was estimated based on the lowest mean cross-validation error (CV-error). To examine genetic differentiation across six localities, analysis of molecular variance (AMOVA) was performed using Arlequin v3.5 [47]. Pairwise *F_ST_* values were also calculated in Arlequin based on allele frequency differences of nuclear SNPs, to estimate difference across populations.

### Statistical analysis

All statistical analyses were performed using R software [48]. We conducted the generalized liner models (GLMs) and linear models (LMs) in the glmmTMB package [49] using two-tailed tests. Model fit and assumptions were assessed using the DHARMa package by inspecting simulated residuals [50].

### Effects of number of subordinates on ecological factors

We analyzed the number of subordinate individuals in relation to ecological factors using GLMs. Because subordinate number was counting data, we fitted models assuming a negative binomial error distribution with a log link function. Shell availability (Shell) varied among territories within localities, whereas predation risk (PR), habitat competition (HC), food competition (FC), and scale-eating predation risk (SE) were measured at the locality level.

Before model construction, we assessed correlations among locality-level predictors. PR and FC were strongly and positively correlated (r = 0.85), whereas all other pairwise correlations were weak (|r| < 0.41). Because PR was the primary ecological variable of interest under the predation-risk hypothesis, we excluded FC from subsequent analyses to avoid overparameterization and facilitate interpretation.

Preliminary model fitting and diagnostic checks using simulated residuals implemented in the R package DHARMa (v. 0.4.7) [50] indicated that global GLMMs including interactions between shell availability and each ecological variable, with locality included as a random effect, produced convergence problems, unstable parameter estimates, and near-zero random-effect variance. Therefore, we removed the random effect from subsequent analyses and used GLMs instead.

To identify which ecological factor modified the relationship between shell availability and subordinate number, we compared an additive baseline model with three candidate interaction models. The baseline model included Shell, PR, SE, and HC as additive fixed effects. Each interaction model included one additional two-way interaction between Shell and one ecological variable: Shell × PR, Shell × SE, or Shell × HC. Thus, the candidate models were subordinate number ∼ Shell + PR + SE + HC, subordinate number ∼ Shell × PR + SE + HC, subordinate number ∼ Shell × SE + PR + HC, and subordinate number ∼ Shell × HC + PR + SE. We first compared each of the three candidate interaction models with the baseline model using likelihood-ratio tests (LRTs) and then evaluated improvement in model fit based on Akaike’s information criterion (AIC).

Model comparisons showed that the Shell × PR interaction substantially improved model fit relative to the additive baseline model (LRT: χ² = 21.21, p < 0.001; AIC reduction = 19.2). The Shell × SE interaction received only weak support (LRT: χ² = 3.89, p = 0.049; AIC reduction = 1.89), whereas the Shell × HC interaction did not improve model fit (LRT: χ² = 0.30, p = 0.59; AIC increase = 1.70). Based on these results, we retained the Shell × PR interaction model as the final model.

### Locality-level differences in body size of group menbers

We analyzed differences in standard length (SL) among localities using a linear model (LM) with locality as a fixed effect, followed by Tukey’s HSD post hoc tests.

### Behaviors, predation risk, and shell availability

To examine how predation risk and shell availability affected the behaviors of breeding females and subordinates, we performed GLM with negative binomial distribution. In these analyses, each behavioral trait was used as a response variable, and we compared a model including the interaction term between Shell and PR (behavior ∼ Shell × PR + SE + HC) with a baseline model without the interaction term (behavior ∼ Shell + PR + SE + HC) using LRTs and AIC values. The final model was then selected based on these comparisons.

As a result, the Shell × PR interaction model was retained for subordinate submissive behavior, subordinate territory maintenance behavior, and breeding female territory maintenance behaviors, whereas the additive model without the interaction term was retained for subordinate aggressive behaviors and breeding female aggressive behaviors.

### Effects of subordinates on breeder behaviors

To test whether subordinates reduce the behavioral workload of breeding female, we performed analyses using negative binomial GLMs. In these modes, the frequency of aggression or maintenance behavior was fitted as response tern of GLMs, and social organization (with or without subordinates) and localities were fitted as fixed terms.

### Effects of shell availability on dispersal from the nest

To examine how experimental shell addition affected changes in juvenile number within natal territories, we analyzed juvenile number in each breeding territory over the experimental period. For each territory, we calculated the change in juvenile number as the number of juveniles 10 days after manipulation minus the number of juveniles on the day of manipulation. We first compared changes in juvenile numbers between treatments using an LM. In addition, to evaluate the direct effect of shell augmentation, we tested the relationship between the change in juvenile numbers and the increase in the number of shells in the nest using a separate LM.

### Effects of experimental removal of subordinates on shell availability

To examine whether shell availability changed following subordinates’ removal, we compared the number of available shells within each nest between 1-day (immediately after removal) and 4-day (four days after removal). Because measurements were paired within the same nests and sample size was small, we used a two-sided Wilcoxon signed-rank test.

## Results

### Social complexity and macro- and micro-ecological factors

Predation risk, the abundance of habitat competitors and food competitors, and shell availability varied substantially among localities (figure 2a–e). The number of subordinates in breeding groups was strongly influenced by the interaction between shell availability and predation risk (GLM: z = 3.803, p < 0.001; figure 2a). In localities with low predation risk, variation in the number of shells had little influence on the number of subordinates. In contrast, in localities with higher predation risk, the number of shells was positively correlated with the number of subordinates. Habitat competition and scale-eating predation risk had no significant effects on the number of subordinates (GLM: z = −0.453, p > 0.50; z = −0.244, p > 0.50).

**Figure 2.**
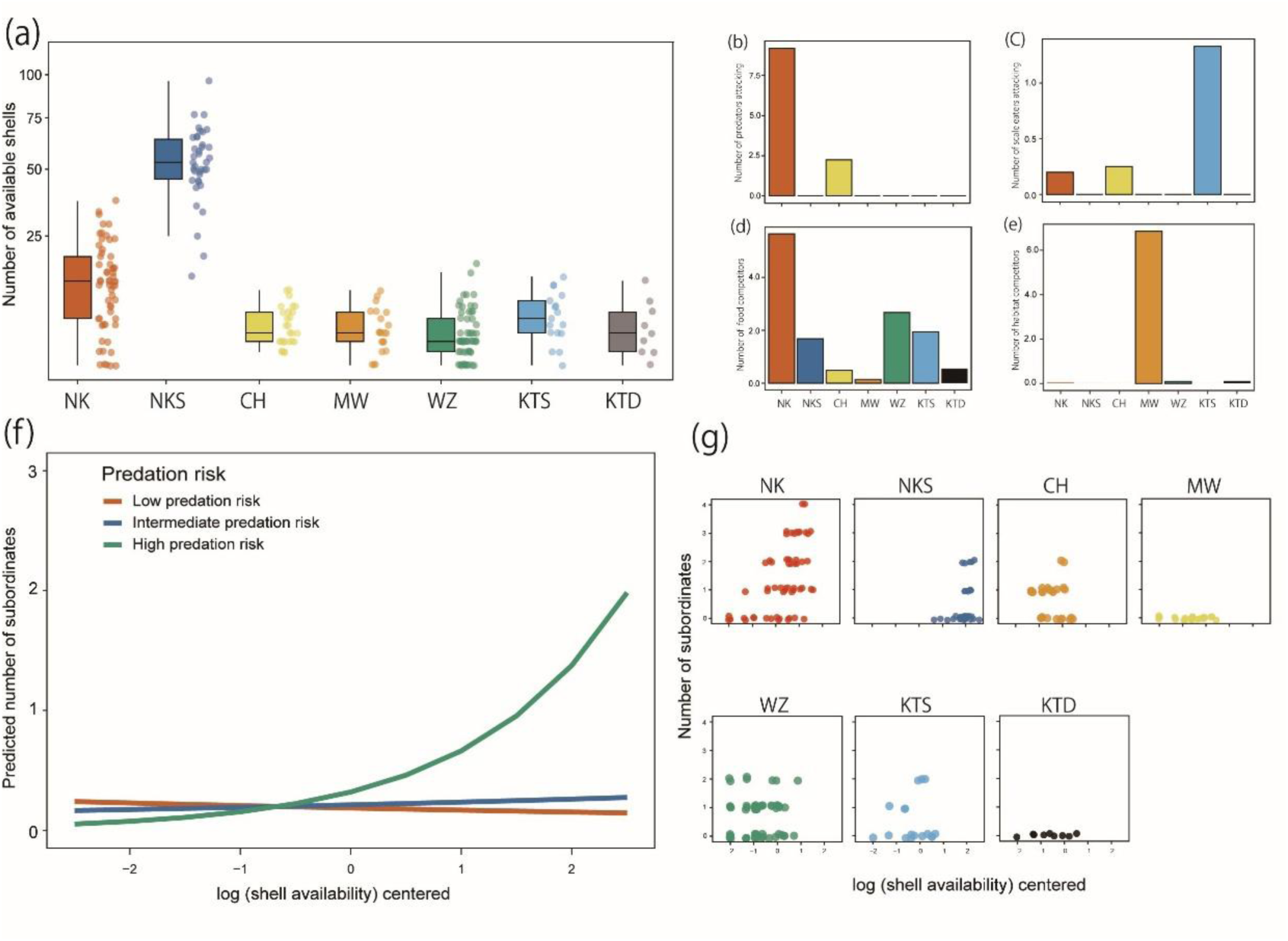
Variation in macro- and micro-ecological factors among study sites and the interactive effects of predation risk and shell availability on the number of subordinates in *Neolamprologus meeli*. (a) Variation in shell availability, a microecological factor, among localities. (b–e) Variation in macroecological factors among localities: (b) number of attacks by predators, (c) number of attacks by scale-eating fishes, (d) number of food competitors observed per transect, and (e) number of habitat competitors around each focal nest. In (a–e), the x-axis indicates locality, and bars represent mean values. (f) Predicted number of subordinates as a function of shell availability and predation risk. Shell availability was log-transformed and mean-centered. Predation risk was fixed at three representative centered values corresponding to NK (high predation risk), CH (intermediate predation risk), and localities where no predators were observed (low predation risk; see figure 2b and table S1). Lines represent predictions from a negative binomial model that included the interaction between shell availability and predation risk. (g) Relationship between shell availability and the number of subordinates in focal territories at each locality. Shell availability was log-transformed and mean-centered. At Nkumbula Island, two habitats were surveyed: a sandy-muddy bottom site (NK) and a shell-bed site (NKS). At Katoto Point, surveys were conducted at a shallow site (KTS; depth < 10 m) and a deep site (KTD; depth > 15 m). The other localities were Wonzye (WZ), Mwina (MW), and Chikonde (CH).

### Difference between body sizes across localities

The standard length (SL) differed significantly among localities in both breeding females and subordinates. Among breeding females, SL varied markedly among localities (one-way ANOVA: F₆,₂₁₇ = 15.16, p < 0.001). Tukey’s HSD post hoc tests revealed that breeding females in KTD point were significantly smaller than those in all other localities. In contrast, breeding females in WZ point and KTS point were significantly larger than those in several other localities (table 2). Among subordinates, SL also differed among localities (one-way ANOVA: F₄,₉₀ = 3.387, p = 0.0125), although this variation was less pronounced than in breeding females. Post hoc comparisons indicated that subordinates in WZ point were significantly larger than those in CH point and NK point, whereas no significant differences were detected among the remaining localities (Table 3).

**Table 3.** Pairwise *F_ST_* among six *Neolamprologus meeli* localities.

| locality | 1 | 2 | 3 | 4 | 5 | 6 |
| --- | --- | --- | --- | --- | --- | --- |
| 1.NK |  |  |  |  |  |  |
| 2.WZ | 0.01983 |  |  |  |  |  |
| 3.CH | 0.0347 | 0.0267 |  |  |  |  |
| 4.MW | 0.03192 | 0.03453 | 0.03975 |  |  |  |
| 5.KTS | 0.13273 | 0.14731 | 0.15851 | 0.14805 |  |  |
| 6.KTD | 0.06541 | 0.07909 | 0.08562 | 0.0938 | 0.05039 |  |

### Effects of shell availability and predation risk on behaviors

Among subordinates, the frequency of submissive behavior toward breeders was strongly influenced by the interaction between shell availability and predation risk (GLM: z = 3.401, p < 0.001). Submissive behavior increased with shell availability under high predation risk, whereas no clear relationship was observed under intermediate or low predation risk (figure 3a). Its frequency was also negatively associated with scale-eating predation risk (GLM: z = −2.190, p = 0.029) but was not significantly associated with the density of habitat competitors (GLM: z = −1.586, p = 0.112).

**Figure 3.**
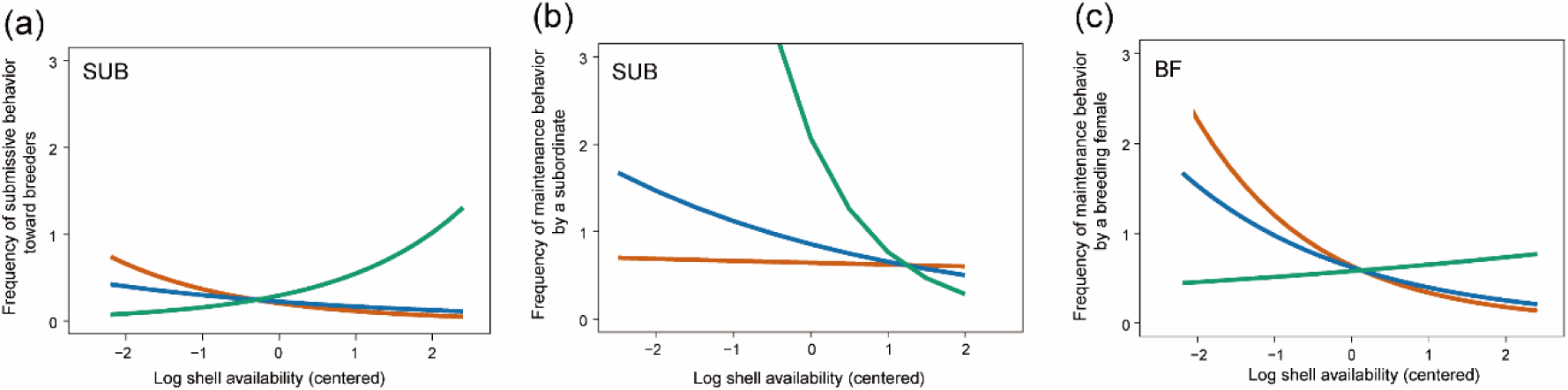
Interactive effects of shell availability and predation risk on the behaviors of subordinates and breeding females. Predicted frequencies of behaviors, which were estimated by GLMs, performed by subordinates and breeding females as a function of shell availability and predation risk. Predictions were obtained from negative binomial generalized linear models. (a) Submissive behaviors by subordinates, (b) maintenance behaviors by subordinates, and (c) maintenance behaviors by breeding females in relation to log-transformed and mean-centered shell availability. Colors indicate predictions at different levels of predation risk. Predation risk was set to three representative centered values, as described for Figure 2f. SUB, subordinates; BF, breeding females.

The frequency of nest-maintenance behavior was also influenced by the interaction between shell availability and predation risk (GLM: z = −2.076, p = 0.038). In contrast to submissive behavior, nest-maintenance behavior decreased with increasing shell availability under high predation risk (figure 3b). Maintenance frequency was positively associated with the density of habitat competitors (GLM: z = 2.135, p = 0.033) but was not significantly associated with scale-eating predation risk (GLM: z = −0.178, p = 0.859).

The frequency of subordinate aggression toward intruders was positively associated with predation risk (GLM: z = 2.219, p = 0.026) and negatively associated with scale-eating predation risk (GLM: z = −2.840, p = 0.006). However, aggression frequency was not significantly associated with shell availability within the nest (GLM: z = −1.379, p = 0.168).

Among breeding females, the frequency of maintenance behavior was influenced by the interaction between predation risk and shell availability (GLM: z = 2.335, p = 0.020). Under low predation risk, breeding females performed nest maintenance more frequently when shell availability was low (figure 3c). Maintenance frequency was also positively associated with scale-eating predation risk (GLM: z = 2.064, p = 0.039) and negatively associated with the density of habitat competitors (GLM: z = −2.008, p = 0.047).

The frequency of aggression by breeding females toward intruders increased with the density of habitat competitors (GLM: z = 3.900, p < 0.001) and decreased with predation risk (GLM: z = −2.921, p = 0.003). However, aggression frequency was not significantly associated with shell availability (GLM: z = 1.577, p = 0.115) or scale-eating predation risk (GLM: z = −1.218, p = 0.223).

### Behaviors of breeding female in social organization across localities

After accounting for locality, the frequency of aggressive behavior by breeding females did not differ significantly between territories with and without subordinates (negative binomial GLM: z = −0.167, p = 0.868). Relative to CH point, aggressive behavior was significantly more frequent at MW point (z = 4.741, p < 0.001), NKS point (z = 3.710, p < 0.001), and WZ point (z = 2.390, p = 0.016). No significant differences from CH point were detected at KTD point (z = 0.720, p = 0.472), KTS point (z = 1.291, p = 0.197), or NK point (z = 1.291, p = 0.197).

The frequency of maintenance behavior by breeding females tended to be more frequent in territories without subordinates, although the difference was not statistically significant (z = 1.740, p = 0.082). Relative to CH point, nest-maintenance behavior was significantly more frequent at KTS point (z = 2.573, p = 0.010) and WZ point (z = 2.354, p = 0.019), but significantly less frequent at NKS point (z = −3.276, p = 0.001). No significant differences from CH point were detected at KTD point (z = 1.867, p = 0.062) or NK (z = 1.382, p = 0.167). The effect for MW point could not be reliably estimated because of the extremely large standard error.

### Effects of shell availability on dispersal from the nest

In the experimentally manipulation for shell availability, the changing rate in juvenile number differed significantly between the experimental and control groups (LMM: t = 3.966, p < 0.001; figure 5a). In the experimental group, in which shells were added, the number of juveniles 10 days after manipulation ranged from -33.3 to 100.0 % of the initial number of juveniles. In contrast, it ranged from -66.7 to 0 % in the control group.

Interestingly, juvenile numbers increased during the 10-day period in three territories in the experimental group. Furthermore, the number of shells was significantly and positively associated with the number of juveniles present in the nests 10 days after manipulation (LMM: t = 3.657, p < 0.005; figure 4b).

**Figure 4.**
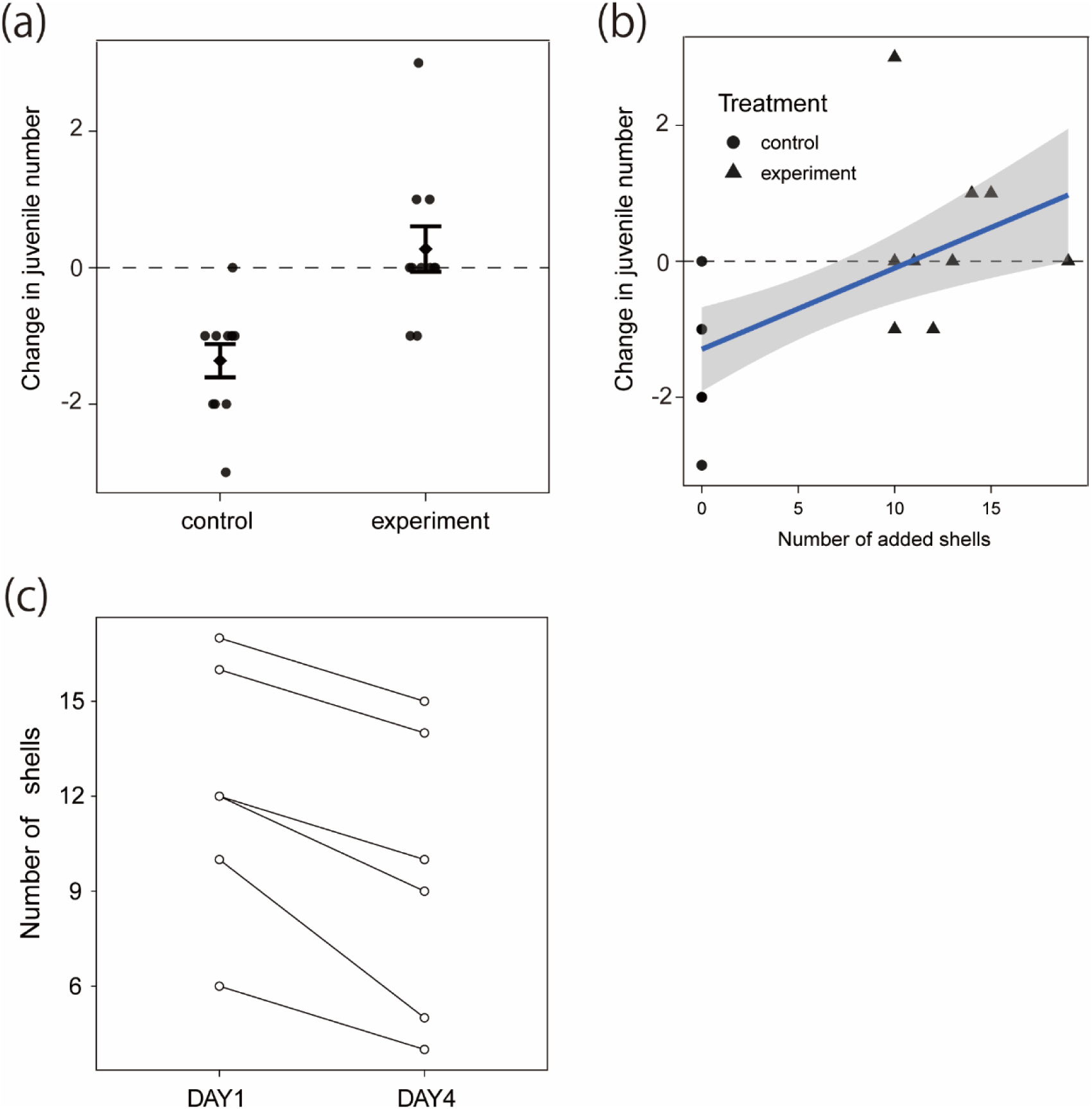
Effects of experimental manipulations for shell availability and subordinates. (a) Change in the number of juveniles within breeding-female territories in the control and shell-addition treatments (n = 22 territories). Points represent individual territories, and error bars indicate the mean ± SE. (b) Relationship between the change in juvenile number and the final number of available shells within breeding-female territories. Circles and triangles represent control and shell-addition territories, respectively. The solid line represents the fitted linear regression, and the shaded area indicates its 95% confidence interval. Positive and negative values indicate increases and decreases in juvenile number, respectively. (c) Change in shell availability following subordinate removal. Each line connects repeated observations of the same nest. Shell availability decreased in all nests between days 1 and 4.

### Effects of experimental removal of subordinates on shell availability

Following subordinate removal, the number of available shells decreased in all focal nests between immediately after removal and four days later (median change = −2.5 shells; Wilcoxon signed-rank test: V = 0, p = 0.031; figure 4c).

## Genetic analysis

Pairwise *F_ST_* values based on 918 SNPs were significantly greater than zero for almost all localities’ pairs (0.020-0.159; permutation test, p < 0.01; table 3), indicating weak but significant genetic differentiation among localities. In the ADMIXTURE analysis, the lowest CV error, indicating the optimal number of clusters, was observed at K = 1, although K = 2 showed a comparably small error (figure S1). Principal component analysis (PCA) revealed partial geographic structuring, with KTS and KTD tending to separate from the eastern populations (NK, CH, MW, and WZ) along PC1, while MW was further differentiated from NK, WZ, and CH along PC2 (figure 5b). Consistent with this pattern, the ADMIXTURE analysis at K = 2 suggested a broad east-west differentiation between Genetic differentiation was relatively weak among localities (figure 5a). In the admixture analysis, the lowest CV error, indicating the optimal number of clusters, was observed at K=1; however, K=2 showed comparably small errors (figure S1). Consistent with this pattern, the ADMIXTURE analysis at K=2 suggested a broad east–west differentiation between the Katoto populations and the remaining populations, despite the absence of discrete genetic clusters (figure 5b).

**Figure 5.**
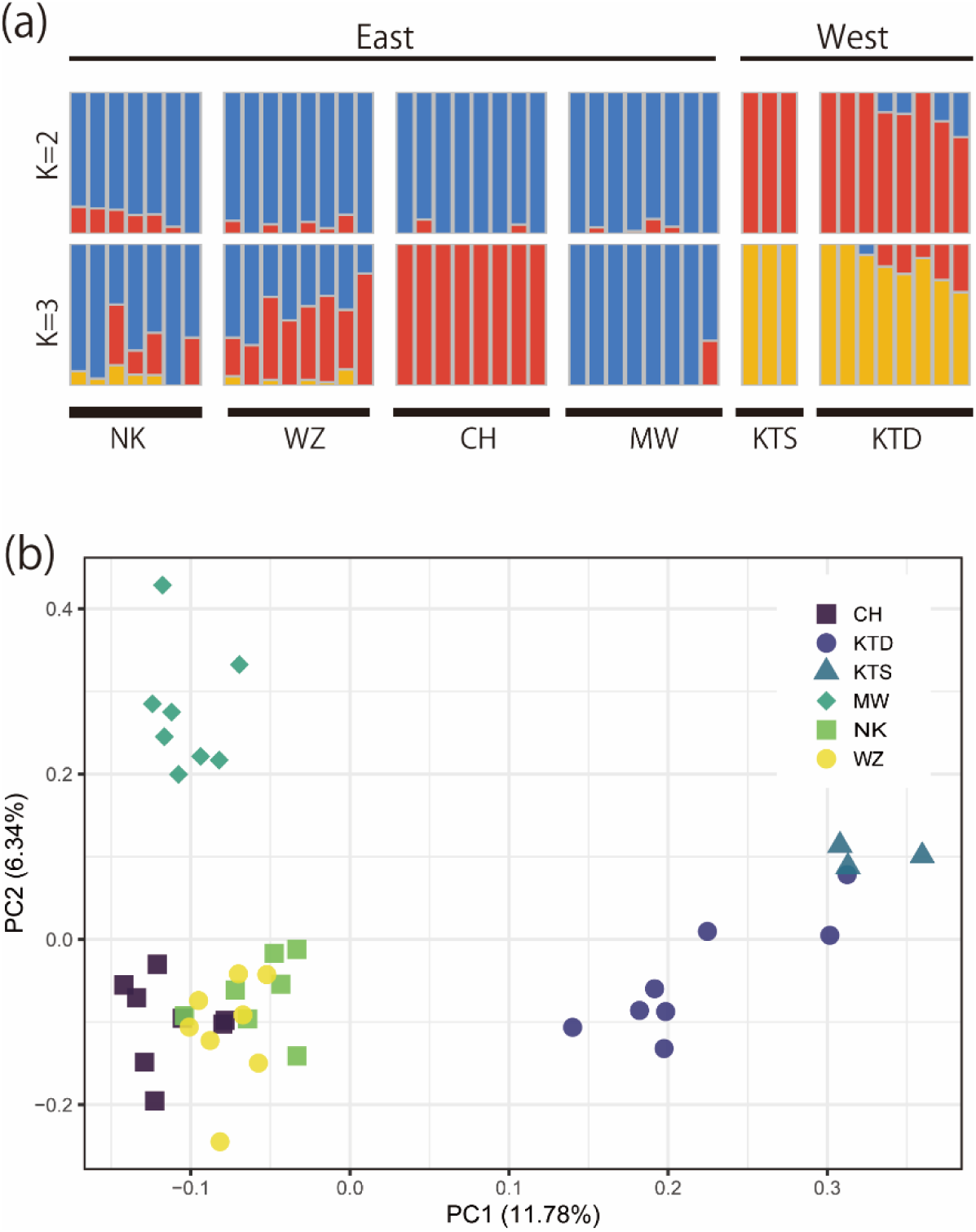
Results of genetic analyses of *Neolamprologus meeli*. These data based on on nuclear SNPs obtained using MIG-seq. (a) ADMIXTURE plots for K = 2 and K = 3. For each value of K, the run with the lowest cross-validation (CV) error among 100 runs is shown. Each vertical bar represents an individual, and the horizontal bars above the plots indicate sampling localities grouped into eastern and western regions. (b) Principal component analysis (PCA). Each point represents an individual, with colors and symbols indicating sampling locality.

## Discussion

In this study, we investigated the ecological factors of social complexity by examining intraspecific variation in social organization across localities of shell-brooding cichlid *Neolamprologus meeli*. We found that social complexity of this species was shaped by the interaction between predation risk and shell availability in their breeding nest. The previous studies on inter- and intra-specific comparison for cichlid’s social complexity suggested that high predation risk may promote delay dispersal and philopatry[7,19,20], which may increase collective action benefits through collective defense and vigilance [24]. Consistent with these studies[7,19,20], we found that the number of subordinate individuals tends to be higher in localities with high predation risk. Furthermore, our results show that even in localities exposed to high predation risk, larger numbers of subordinates could be maintained only when shell availability was high. This suggests that while predation risk increases the collective action benefits, territory quality such as shell availability, constrains the upper limit of group size [51].

Behavioral observations showed that not only nest maintenance and aggression toward intruders by subordinates, but also the frequency of submissive behavior toward breeders, which is unlikely to be directly related to predator defense, varied with the combination of predation risk and shell availability. In particular, under high predation risk, subordinates exhibited submissive behavior toward breeders more frequently when shell availability was high. High predation risk is likely to increase the costs of dispersal [13,19,20], whereas abundant shell resources may increase the benefits of remaining in the current territory. Under these conditions, subordinates may therefore invest more heavily in appeasement behavior, thereby increasing breeders’ tolerance of their continued residence or delayed dispersal. This interpretation is consistent with the pay-to-stay hypothesis [28,30] , which proposes that helping by subordinates functions as a form of appeasement that allows them to remain in the group [28,31]. The greater frequency of subordinate aggression toward intruders under high predation risk may similarly serve an appeasement function [28,31]. However, such aggression may also provide direct or indirect fitness benefits by protecting the subordinates themselves or their relatives. Moreover, variation in aggression frequency may reflect trade-offs in time allocation between defense and nest maintenance [52,53] or the division of helping tasks among group members [34,54,55].

Our shell-addition experiment conducted in a high-predation-risk environment provided direct evidence that shell availability influences juvenile retention and recruitment. Increasing shell availability led to an increase in the number of juveniles within breeder territories, and in some cases juvenile numbers exceeded their pre-manipulation levels. In *N. meeli*, intense sibling competition over shells occurs early in development: individuals excluded from the nest are forced to disperse, whereas those that remain may later become subordinate helpers [34]. During the experiment, we also observed juveniles entering breeder territories shortly after shells were added (YY, personal observation). Although we could not determine whether these individuals had previously been excluded from the same territories or had immigrated from elsewhere, these observations suggest that juveniles respond rapidly to changes in nesting-resource availability. Together with the observational results, the experiment supports the hypothesis that shell availability constrains juvenile retention and recruitment, and consequently the size and composition of social groups, under high predation risk.

At the same time, the experimental removal of subordinates demonstrated that they contribute to maintaining shell availability within breeding territories. Together, the two experimental manipulations suggest reciprocal feedback between group composition and territory quality. Shell availability facilitates juvenile retention and recruitment, whereas subordinates maintain shell availability by preventing shells from becoming buried. Territory quality is therefore not merely an external ecological factor that determines group size but is also partly generated and maintained through the activities of group members. Consistent with this interpretation, at the locality characterized by high predation risk, subordinates performed nest maintenance more frequently when shell availability was low. This result may suggest that subordinates are more strongly motivated to maintain or increase shell availability under high predation risk. Such subordinate-mediated improving territory quality may generate a positive feedback whereby existing subordinates maintain the capacity of territories to retain juveniles, some of which may subsequently become subordinates themselves.

One potential benefit of helping in cooperative breeders is load-lightening, whereby breeders reduce their own effort in response to contributions by subordinates [51,56]. However, we found no consistent association between subordinate presence and the rates of nest maintenance or aggression toward intruders by breeding females. Thus, our results provide little evidence that subordinates reduce these components of female workload. This does not necessarily mean that subordinate contributions provide no benefits to breeders. If breeding females maintain their own effort, subordinate contributions may be additive, increasing total nest maintenance and defense rather than replacing breeder effort [34]. Consistent with this interpretation, experimental removal of subordinates reduced shell availability, demonstrating that subordinates make a functional contribution to territory maintenance.

Moreover, previous work in *N. meeli* suggests that subordinates can reduce the workload of breeding males [34], indicating that load-lightening may be sex-specific. Subordinates may also provide benefits not captured by our behavioral measures, including enhanced predator detection and improved territory quality. Assessing reproductive success, survival, and future fecundity will therefore be necessary to determine the net fitness consequences of subordinate behavior [56].

Finally, ADMIXTURE and pairwise *F_ST_* analyses of the MIG-seq data revealed no distinct genetic clustering and only weak genetic differentiation among localities. Thus, locality-level variation in social organization is unlikely to be explained by broad-scale population structure alone and is consistent with a substantial influence of local ecological conditions. Nevertheless, weak genome-wide differentiation does not exclude genetic contributions involving a small number of loci or genotype-by-environment interactions. Further studies combining higher-resolution genomic analyses with common-garden or reciprocal-transplant experiments will be necessary to disentangle environmental and genetic contributions to variation of social complexity [57].

In conclusion, our study shows that intraspecific variation in the social organization of *N. meeli* is shaped by the interaction between predation risk and shell availability. Predation risk may increase the benefits of remaining in groups, whereas shell availability influences the capacity of territories to retain and recruit juveniles. Our experiments further reveal a reciprocal relationship between group composition and territory quality: shell availability promotes juvenile retention and recruitment, while subordinates maintain shell availability through nest maintenance. Together with the weak genome-wide differentiation among localities, these findings support an important role for interaction of macro- and micro-ecological factors in generating and maintaining social complexity.

## Ethics statement

All experimental protocols were approved by the Animal Care and Use Committees of Kyoto University and adhered to the ASAB/ABS guidelines for the treatment of animals in behavioral research. Our field research in Lake Tanganyika was conducted with permission from the Zambian Ministry of Agriculture, Food, and Fisheries and complied with the current laws in Zambia.

## AI statement

In preparing this manuscript, we use ChatGPT 4.0 to refine the English text.

## Acknowledgments

The members of the Maneno Tanganyika Research Team, Laboratory of Aniaml Ecology, Kyoto University for Advanced Studies provided helpful comments and discussion. This study was financially supported by JST SPRING (A9425060500043, A9426060500043), DoGs [to Y.Y.] and Hakubi project of Kyoto University, and JSPS KAKENHI (25K02309, 23KK0131, and 23H038689) [to S.S.]

## Author contribution

Experimental design: Y.Y., S.S. Field observation and sampling: Y.Y., S.S., R.H., R.I. Collecting behavioral data: Y.Y., S.S. Collecting predation risk data: Y.Y., K.K. field manipulative experiment: Y.Y., Statical analysis: Y.Y. Support of statical analysis: S.S. genetic analysis: Y.Y., Y.T. Writing of the manuscript: Y.Y., S.S.

## Data availability

The data and R script for analysis that support the findings of this study are available in Dryad (https://XXXXXX)

## Supplementary figures

**Fig.S1.**
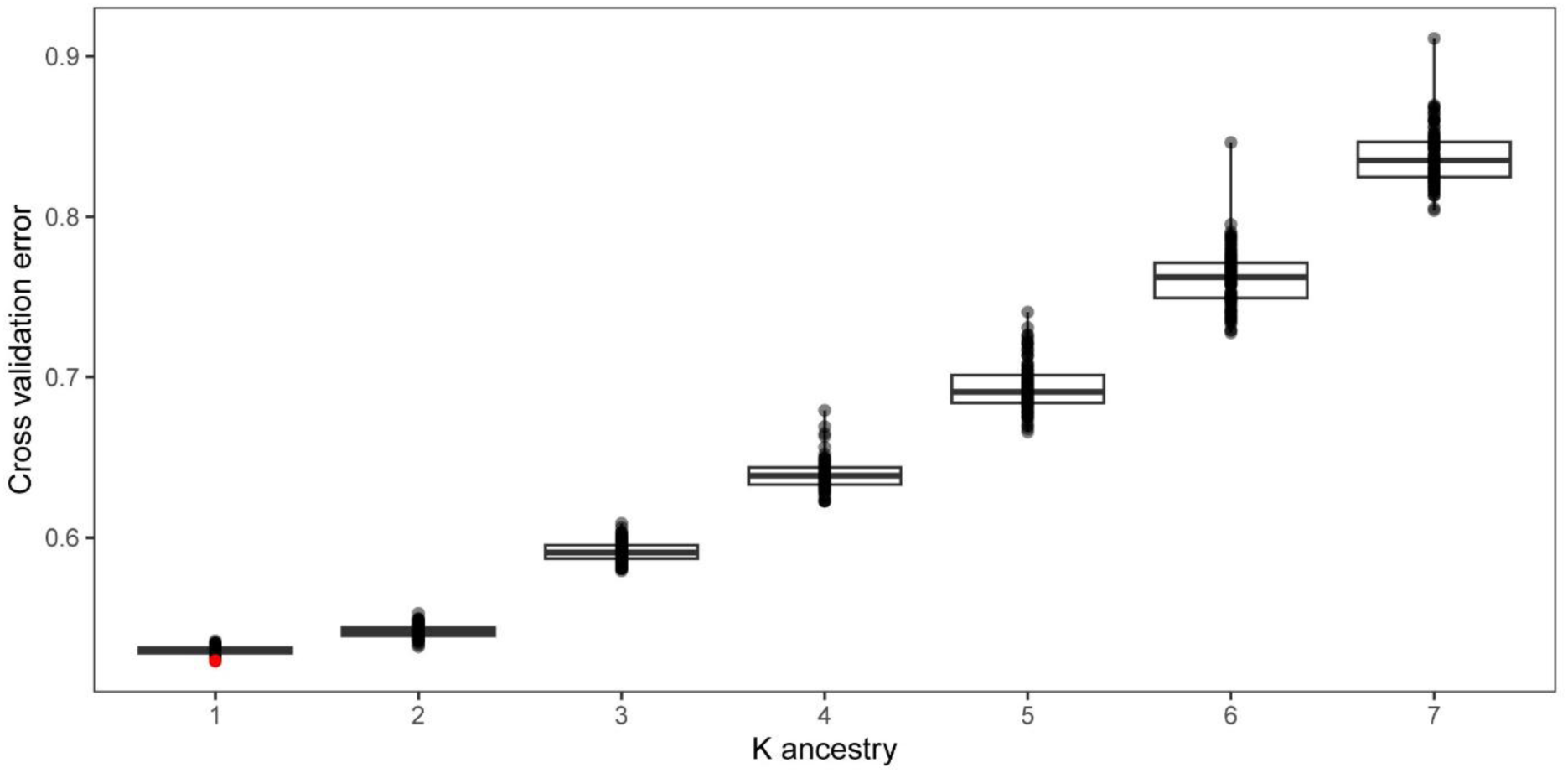
Cross validation (CV) error of 100 trials for each K in ADMIXTURE analysis

**Table S1.**
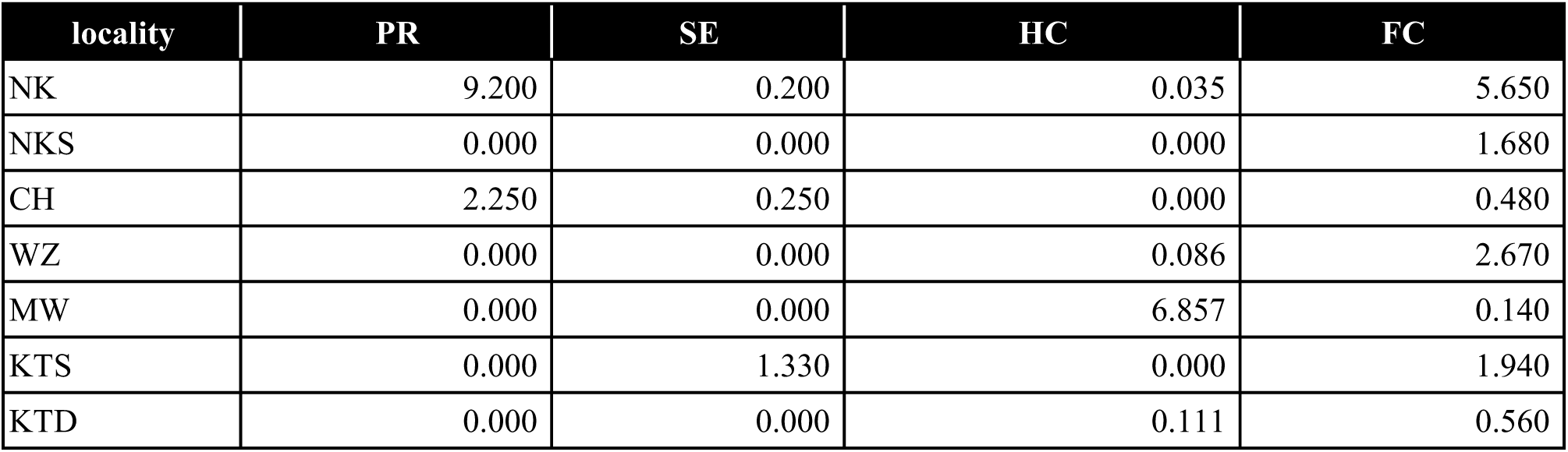
Data of ecological factors for each locality. These values are average of ecological factors for each locality. (PR: predation risk, SE: scale-eating predation risk, HC: habitat competition, FC: Food competition)

